# Tafenoquine inhibits replication of SARS-Cov-2 at pharmacologically relevant concentrations *in vitro*

**DOI:** 10.1101/2020.07.12.199059

**Authors:** Geoffrey S. Dow, Angela Luttick, Jen Fenner, David Wesche, Karen Rowland Yeo, Craig Rayner

## Abstract

Tafenoquine [TQ] exhibited EC50/90s of ~ 2.6/5.1 μM against SARS-CoV-2 in VERO E6 cells and was 4-fold more potent than hydroxychloroquine [HCQ]. Time-of-addition experiments were consistent with a different mechanism for TQ v HCQ. Physiologically based pharmacokinetic (PBPK) modeling suggested that lung unbound concentrations of TQ in COVID-19 patients may exceed the EC90 for at least 8 weeks after administration. The therapeutic potential for TQ in management of COVID-19 should be further evaluated.

## MANUSCRIPT TEXT

Therapeutic options with conditional or emergency approval by regulators for hospitalized COVID-19 patients include dexamethasone and remdesivir [8–9, 21]. However, although more than a 1000 clinical trials are underway to evaluate other alternatives, there are not yet any approved options for prevention [pre-exposure or post-exposure prophylaxis (PrEP or PEP)] of COVID-19 or early treatment or treatment of mild COVID −19 infection in either out-patient or hospitalized settings. Tafenoquine [TQ], an FDA-approved 8-aminoquinoline [8AQ] antimalarial, can be administered orally, weekly, for up to six months, and is differentiated relative to the 4-aminoquinolines [4AQs] hydroxychloroquine (HCQ) and chloroquine (CQ) in that it does not prolong QTc interval and acts more broadly against a variety of malaria life cycle stages via different mechanisms of action [1, 6, 10]. Specifically, 4AQs kill malaria parasites with the same mode of action as has been proposed for their anti-coronavirus effects, by increasing intravacuolar pH [22, 24]. While mechanisms of action of 8AQs are poorly defined, recent studies have suggested malaria parasites are killed site-specifically by hydrogen peroxide accumulation generated by a two-step biochemical relay involving monooxygenases and cytochrome P450:NADPH oxidoreductase [6]. Also, tafenoquine is active against the lung pathogen *Pneumocystis in vitro* and *in vivo* at pharmacologically relevant concentrations and doses [4, 16]. Given the dearth of therapeutic options for COVID-19, we sought to explore the pharmacological plausibility and proof of hope [12] for TQ as a potential therapeutic candidate for further evaluation in the prevention and early treatment phase of COVID-19.

We first evaluated the antiviral activity of TQ against SARS-CoV-2-Australia/VIC01/2020 [a gift from the Doherty Institute, Melbourne, Australia] in a cytopathic effect [CPE] assay. Briefly, SARS-CoV-2-infected VERO E6 cell monolayers [at multiplicity of infection [moi] of 0.05] in 96-well plates were incubated at 37°C in 5% CO_2_ while exposed to differing concentrations of TQ, remdesivir [RD] and HCQ in triplicate wells in minimum essential medium supplemented with 1% (w/v) L-glutamine, 2% fetal bovine serum and 0.2% DMSO [the latter for compound dissolution]. [The same medium was used for other viral assays described later]. After 4 days, cell viability was assessed via MTT and the EC50 and CC50 calculated using the method of Pauwels *et al* [15]. TQ, HCQ, and RD exhibited EC50s of 15.6, > 100, and 0.77 μM respectively, and TQ exhibited greater selectivity than HCQ [selectivity index of 2.4 versus < 1.0, respectively, see Table 1]. An EC50 could not be established for HCQ as the CC50 was lower than the highest concentration tested [see Table 1]. Thus, the validity of the test system used was confirmed by virtue of the observation of a sub-micromolar EC50 for RD, and TQ was found to exhibit more potent antiviral activity than HCQ.

**Table 1:** EC_50_, CC_50_, and selectivity data.

|  | 4-Day CPE Assay |  |  | 2-Day YR Assay |  |  |
| --- | --- | --- | --- | --- | --- | --- |
|  | EC<sub>50</sub><br>[μM] | MTT CC<sub>50</sub><br>[μM]* | SI | EC<sub>50</sub><br>[μM] | EC<sub>90</sub><br>[μM]** | SI*** |
| Tafenoquine | 15.7 | 37.2 | 2.4 | 2.6 | ~5.1 | 14.2 |
| HCQ | 43%<br>inhibition at<br>100 μM | 55.5 | NA | 10.4 | ~21 | 5.3 |
| Remdesivir | 0.77 | NT | NA | NT | NT | NA |

*\*CC<sub>50</sub> for 4-Day CPE assay determined using MTT \*\*Determined by multiplying EC<sub>50</sub> by 1.98 [19] \*\*\* For the 48h yield reduction assay selectivity was determined with respect to CC<sub>50</sub> calculated from the 4-day MTT assay. NT = not tested. NA =not applicable*

TQ’s ability to interfere with infectious virus replication and reduce the yield of progeny virus was assessed using a yield reduction [YR] assay. Briefly, SARS-CoV-2-infected VERO E6 cells [moi 0.05] in 24-well plates were incubated at 37°C in 5% CO_2_ while exposed to differing concentrations of TQ and HCQ. After 48h, supernatant was harvested and added to 96-well plates in triplicate, incubated for three days at 37°C in a humidified 5% CO_2_ atmosphere, after which virus-induced CPE scored visually by two independent operators. The TCID50 of the virus suspension was determined using the method of Reed-Muench [17]. EC50s, calculated using the method Pauwels et al [15] were 2.6 and 10.4 μM for TQ and HCQ respectively [Table 1]. EC90s, estimated by multiplying the EC50 by 1.98 [19], were 5.1 and 21 μM respectively [Table 1] We note that the EC50 was reduced to 2.6 μM from 15.6 μM when virus is exposed to TQ for 48 v 96h, suggesting that the anti-viral effect of TQ is dependent on viral load.

A time-of-addition [TOA] assay was used to identify when TQ interferes with the viral replication cycle and to investigate the mechanism of action. A compound that acts early in the replication cycle should reduce the virus titers at early time points; conversely, a compound that acts later in the replication cycle should reduce the virus titers at later time points. Antiviral compounds that act early in the cycle may be interfering with the attachment of the virus into the host. Compounds that act in the middle of the cycle may be affecting one or more of the many processes involved in viral replication. Compounds that act late in the cycle may be affecting processes associated with viral release. The TOA experiment was conducted using a variation on the method of Schneider et al [19]. Briefly, the virus was allowed to adsorb to cell monolayers [moi of 1] in 24-well plates for 1h, then parallel cultures were incubated at 37°C in 5% CO_2_ for 15, 30, 60 minutes, 2, 4, 6 hours prior to adding 15μM HCQ, 15μM TQ and 5μM RD, or negative control (assay media only). Concentrations were chosen to be approximately 5-fold higher than the EC50 in the 48h YR assay, except that for HCQ this was not possible due to cytotoxicity at concentrations ≥ 50 μM, so TQ and HCQ were compared head-to-head at the same concentration [15 μM]. For the 0-minute time point, test or control article were added immediately following virus pre-adsorption. Plates were incubated at 37°C in 5% CO_2_ until 8h after viral adsorption [8h representing the duration of one cycle of replication]. Supernatants were harvested and virus titer determined at each time point via virus yield assay as described earlier. TQ reduced virus titers when added to infected cultures between 0 and 4h, whereas HCQ did not exhibit a discernable effect regardless of the time of addition [Figure 2]. This observation provides indirect evidence of a differing mode of action between the 8 and 4-aminoquinolines against SARS-CoV-2 that warrants further investigation.

**Figure 1:**
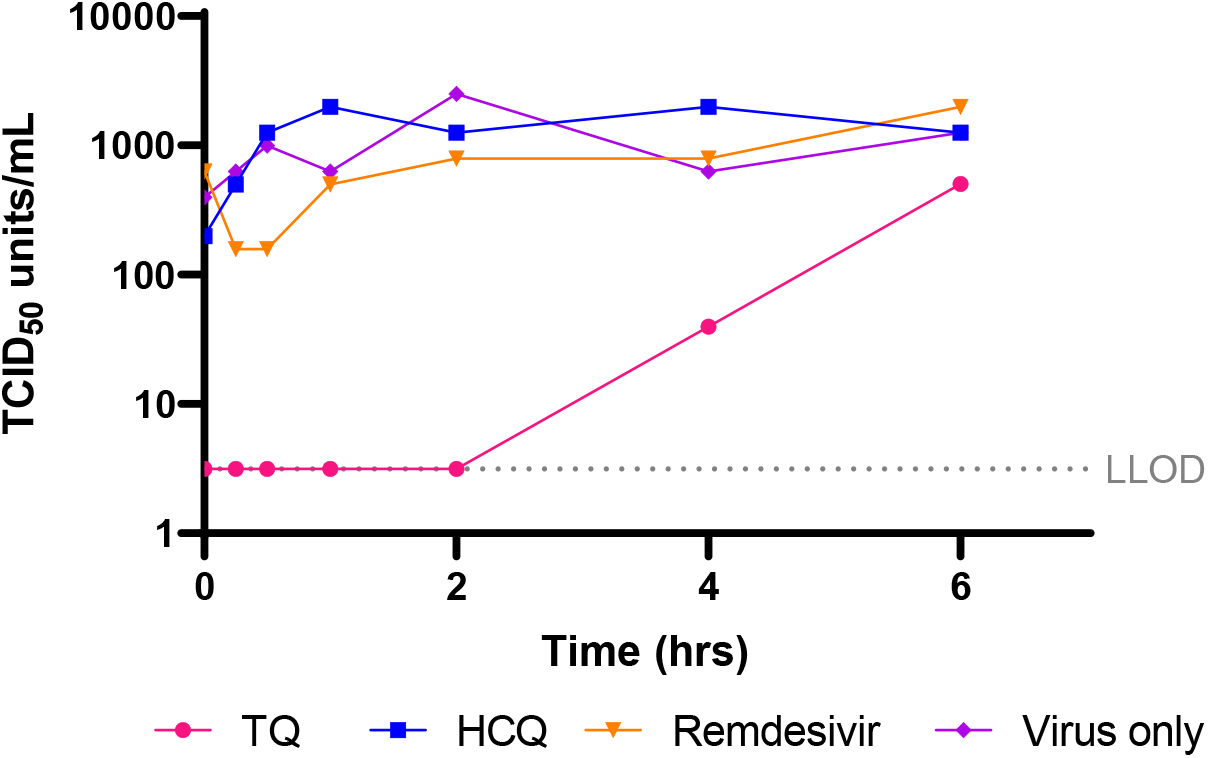
Time of addition experiments. Tafenoquine [15 μM, TQ], hydroxychloroquine [15 μM, HCQ], remdesivir [5 μM], or virus control were added at the indicated time points to Vero cells to which SARS-CoV-2 was allowed to pre-adsorb for 1 h. At 8h virus titer was determined by viral yield assay. Each point on the graph represents the virus titer present after one cycle of replication following addition of drug at the indicated time following virus adsorption.

**Figure 2:**
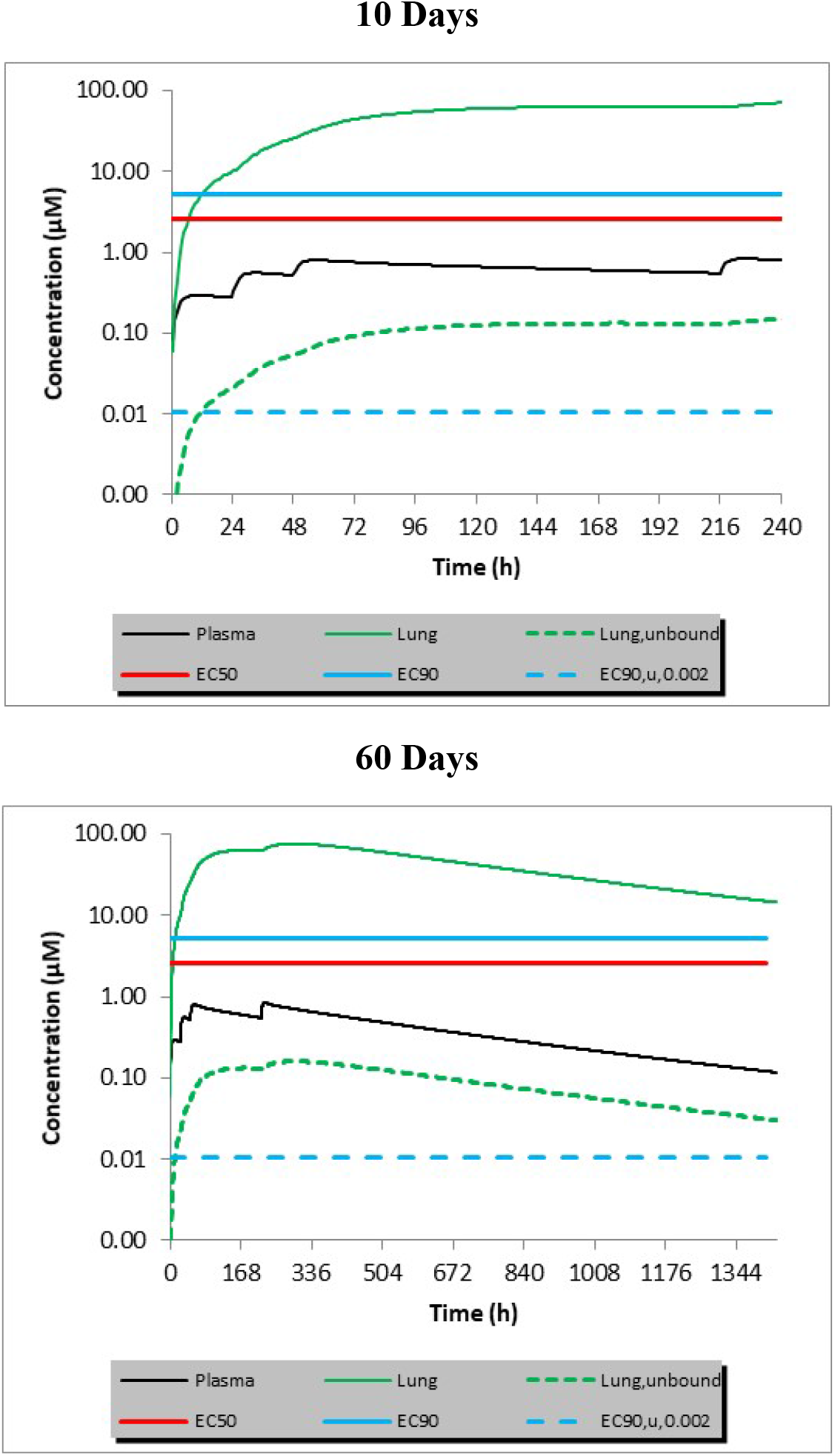
Predicted concentration-time curves for plasma, lung, and lung unbound following administration of tafenoquine at the approved dose for malaria prophylaxis [200 mg/day for three days followed by 200 mg one week later], in which the pH of lung tissue was assumed to 6. The EC50, EC90, and EC90_u_ concentrations from the YR assay are indicated as horizontal lines across the figure.

For the purposes of establishing proof of hope (i.e. expectation that pharmacologically relevant concentrations at the target site can be achieved clinically and a likelihood that a relevant therapeutic index exists), we sought to estimate TQ lung concentrations achieved following the first four doses of the FDA-approved dose of TQ for malaria prophylaxis [200 mg/day for three days, followed by 200 mg seven days later [1]]. Whole lung, lung-unbound, and plasma concentrations at an assumed lung pH of 6 were simulated using Simcyp [V19.1, Simcyp Ltd, Sheffield, UK] as described by Rowland Yeo et al, [18]. A pH of 6 was selected, since slightly acidic pH is anticipated for COVID-19 patients. The input assumptions for various parameters are described in Table 2. The method was first validated against published PK data sets for a single dose of 300 mg, a 1200 mg dose split as equal divided doses over three days, and a single dose of 200 mg [2, 11, 13]. Concentration-time profiles were plotted against the EC50, EC90, and EC90_u_ values from the 48 h YR assay. The EC90_u_ was determined by multiplying the EC90 by the estimated unbound fraction to correct for non-specific binding in culture medium containing 2% FBS, a protein concentration of approximately 90 ng/mL. Assuming that efficacy at the site of action is determined by the unbound drug, predicted lung unbound concentrations exceed the EC90_u_ of tafenoquine for at least 8 weeks following administration of the second dose of the proposed regimen [Figure 2].

**Table 2:**
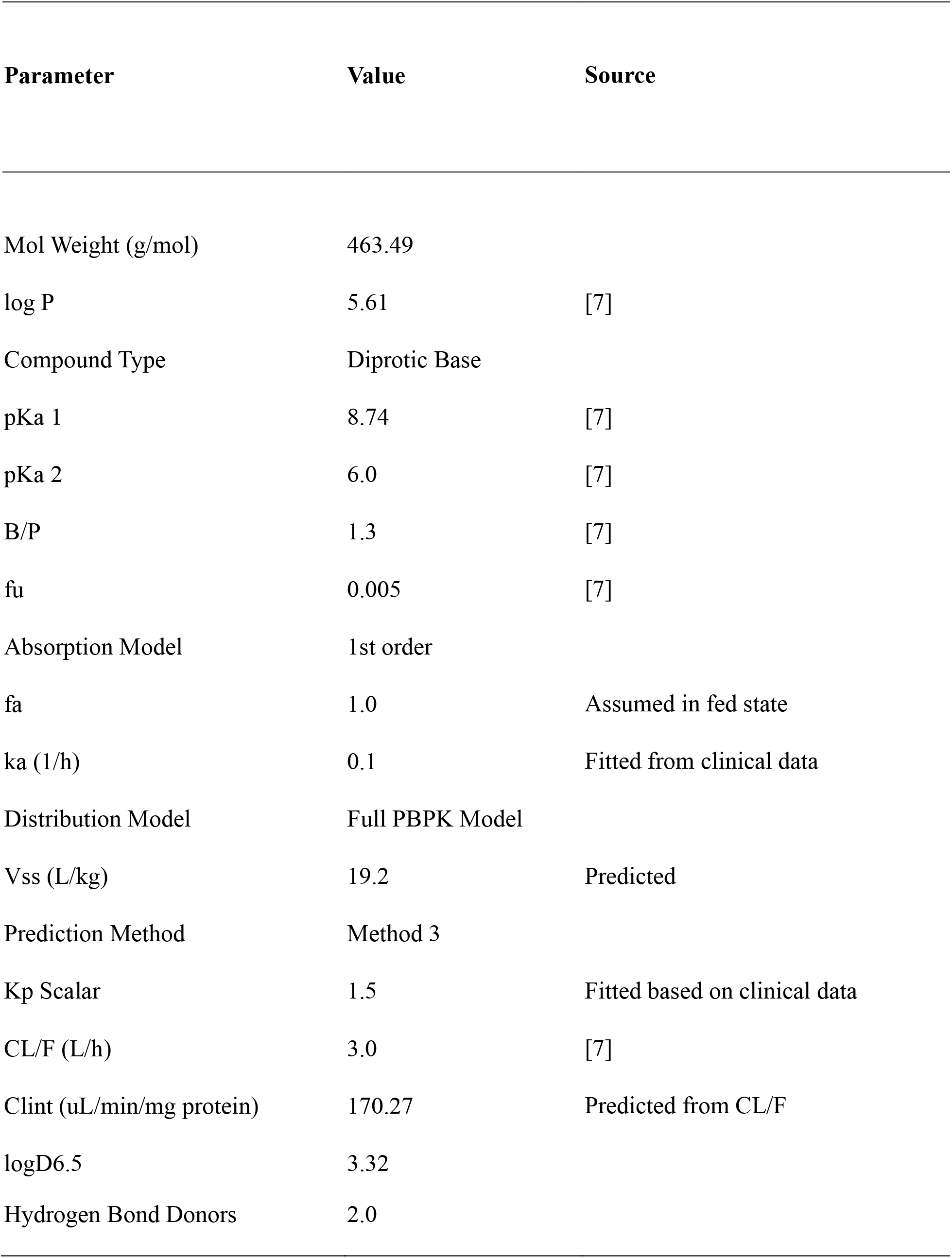
PK modeling assumptions.

The urgency of identifying therapeutic options for COVID-19 patients, and the absence of widely available animal models, has meant that repositioning programs have embraced candidate prioritization through comparison of SARS-CoV-2 *in vitro* susceptibility in VERO E6 cells to anticipated plasma or unbound lung concentrations of proposed or approved clinical dosing regimens [3, 18, 23]. It is apparent that the lack/modest level of activity of HCQ in both animal models and clinical studies [5,14] was not predicted from VERO cells and evidence of extensive lung penetration of HCQ. There is much speculation about the mechanistic basis and generalizability of this incongruence, and a growing consensus on the need to move towards including additional *in vitro* cell systems evaluations when evaluating COVID-19 candidates, such as lung epithelial cells and organoids that may more closely represent the *in vivo* situation. However, in the absence of international guidance and harmonization on appropriate *in vitro* cell systems to evaluate antiviral activity of SARS-CoV-2, VERO cells will continue to be the standard.

We have demonstrated antiviral activity of TQ against SARS-CoV-2 in *in vitro* VERO E6 cell system at concentrations deemed to be pharmacologically relevant and achievable in lung tissue via PBPK modelling at doses of TQ which have been approved. Furthermore, we have noted indirect evidence that is suggestive of a differentiated mechanism of action of this 8AQ against SARS-CoV-2. Based on these data, we believe that there is pharmacological plausibility and proof of hope that TQ may have potential to be effective in the treatment pathway for COVID-19, and would encourage further evaluation.

## ACKNOWLEDGEMENTS

GSD is the majority equity holder in 60 Degrees Pharmaceuticals LLC [60P] which has a financial interest in the commercial success of tafenoquine. GSD is an inventor on several tafenoquine-related COVID 19 patents, for which ownership interest has been assigned to 60P. AL and JF are employees of 360 Biolabs and have no personal financial relationship to 60P. CR, KRY, and DW are employees of Certara and have no personal financial relation with 60P or the Gates Foundation. 60P provided funding for antiviral assays performed by 360 Biolabs under contract. The Gates Foundation [BGMF] provided funding for the pharmacokinetic modeling. All authors contributed to generation of the data, and writing of, this manuscript. We are grateful for helpful comments on the manuscript and methods from Bryan Smith, Jennifer Herz and Ty Miller from 60P, Josh Berman and Bill Ravis from Fast Track Drugs and Biologics, and for project coordination efforts by Jennifer Herz from Biointelect Pty Ltd.

